# Bidirectional emotional regulation through prefrontal innervation of the locus coeruleus

**DOI:** 10.1101/2024.02.07.579259

**Authors:** Mayumi Watanabe, Akira Uematsu, Joshua P. Johansen

## Abstract

Traumatic experiences produce powerful emotional memories which can subsequently be enhanced or reduced through cognitive control mechanisms. Noradrenaline from the brainstem locus coeruleus (LC) is activated during aversive emotion-inducing experiences and upregulated in individuals suffering from anxiety and trauma related disorders. The medial prefrontal cortex (mPFC) participates in cognitive and emotional control processes such as production of learned defensive responses and suppression of aversive memories through extinction. However, it is unclear whether or how distinct mPFC regions influence the LC to regulate learned emotional responding. Using viral based anatomical tracing techniques, we found that the LC receives topographically organized projections from the prelimbic (PL) and infralimbic (IL) subregions of mPFC. Furthermore, optogenetic approaches revealed that PL and IL inputs to LC are required to inhibit or facilitate, respectively, the extinction of aversive memories. Moreover, LC-projecting neurons in PL and IL exhibited distinct activity patterns during extinction learning, with IL-to-LC neurons displaying sustained, sensory cue-evoked activation, while activity in PL-to-LC inputs is elevated during periods of externally and internally generated aversive emotional responding. Together, these results demonstrate that mPFC subregions bidirectionally regulate extinction of emotional memories through differential modulation of the LC-noradrenaline system.

## Main Text

Learning to predict threats through experience is essential for survival, but it is also important to extinguish aversive emotional responses when they are no longer appropriate. Deficits in extinction of aversive memories are associated with pathological states such as posttraumatic stress disorder (PTSD). The neuromodulator noradrenaline plays important roles in emotional learning as well as the extinction of emotional responding when it is no longer appropriate^1–6^ and dysfunction of noradrenergic signaling is observed in anxiety- and stress-related disorders^7,8^. The locus coeruleus (LC), the major source of noradrenaline to forebrain, modulates fear, anxiety and extinction through projections to amygdala and prefrontal cortex^9–13^. However, how afferent inputs to the LC-noradrenaline system regulate emotional processing is poorly understood.

The medial prefrontal cortex (mPFC) is an important brain region for context dependent cognitive processing and emotion. mPFC neurons project to the LC^14–16^ and a neural circuit from mPFC to LC has been hypothesized to provide top-down control of the LC-noradrenaline system as an interface between cognitive cortical systems and central noradrenaline function. Specifically, early theoretical work proposed that prefrontal inputs to LC provide cognitive regulation to enable flexible behavior^17^. Moreover, electrophysiological studies demonstrated functional connectivity from mPFC to LC^18–21^ and recent studies reported that prefrontal inputs to the LC mediate control of arousal^15^ and pain modulation^22^. In regulation of aversive emotional responses, prelimbic (PL) and infralimbic (IL) subregions of mPFC have complementary roles; PL neurons drive expression of learned fear responses and IL activity promotes formation of extinction memories. This suggests that PL and IL inputs to LC may differentially influence fear extinction, but it has been unclear whether and how inputs from prefrontal subregions to LC participate in fear extinction.

Using viral tracing and functional manipulation approaches, we show that the LC receives topographically organized projections from PL and IL and that PL and IL inputs to LC opposingly regulate the formation of fear extinction memories. Furthermore, these separate mPFC-to-LC pathways exhibit distinct activity patterns during fear extinction learning, with PL activity being related to internally and sensory driven aversive emotional responses and IL neurons providing more sustained responding through the extinction session. These results suggest that the mPFC provides top-down regulation of the LC-noradrenergic system for adaptive control of emotional responding, with distinct mPFC subregions constraining or facilitating extinction memory formation.

## Results

### Anatomical connectivity from prefrontal subregions to the locus coeruleus

We first examined whether PL and IL subregions of mPFC project to LC. To study this question, we injected Retrobeads into LC of wild-type rats and found labeled neurons in both PL and IL (***Fig. 1a***, *n*=3). To examine whether LC-noradrenergic neurons receive direct monosynaptic inputs from PL and IL, we used trans-synaptic rabies virus tracing. We injected adeno-associated viruses (AAV) carrying TVA and glycoprotein under the control of Cre-recombinase into the LC of tyrosine hydroxylase (TH)-Cre recombinase rats^23^. Three weeks after AAV injection, we injected the modified CVS rabies virus strain^24^ carrying green fluorescent protein (GFP), allowing for transsynaptic infection of synaptic inputs onto LC-noradrenergic neurons. Starter cells expressing both glycoprotein and GFP were found in LC-noradrenergic neurons (***Fig. 1b***) and we identified GFP-expressing input cells in both PL and IL regions (***Fig. 1c***, *n*=2). These results demonstrate that LC-noradrenergic neurons receive direct monosynaptic inputs from PL and IL.

**Figure 1.**
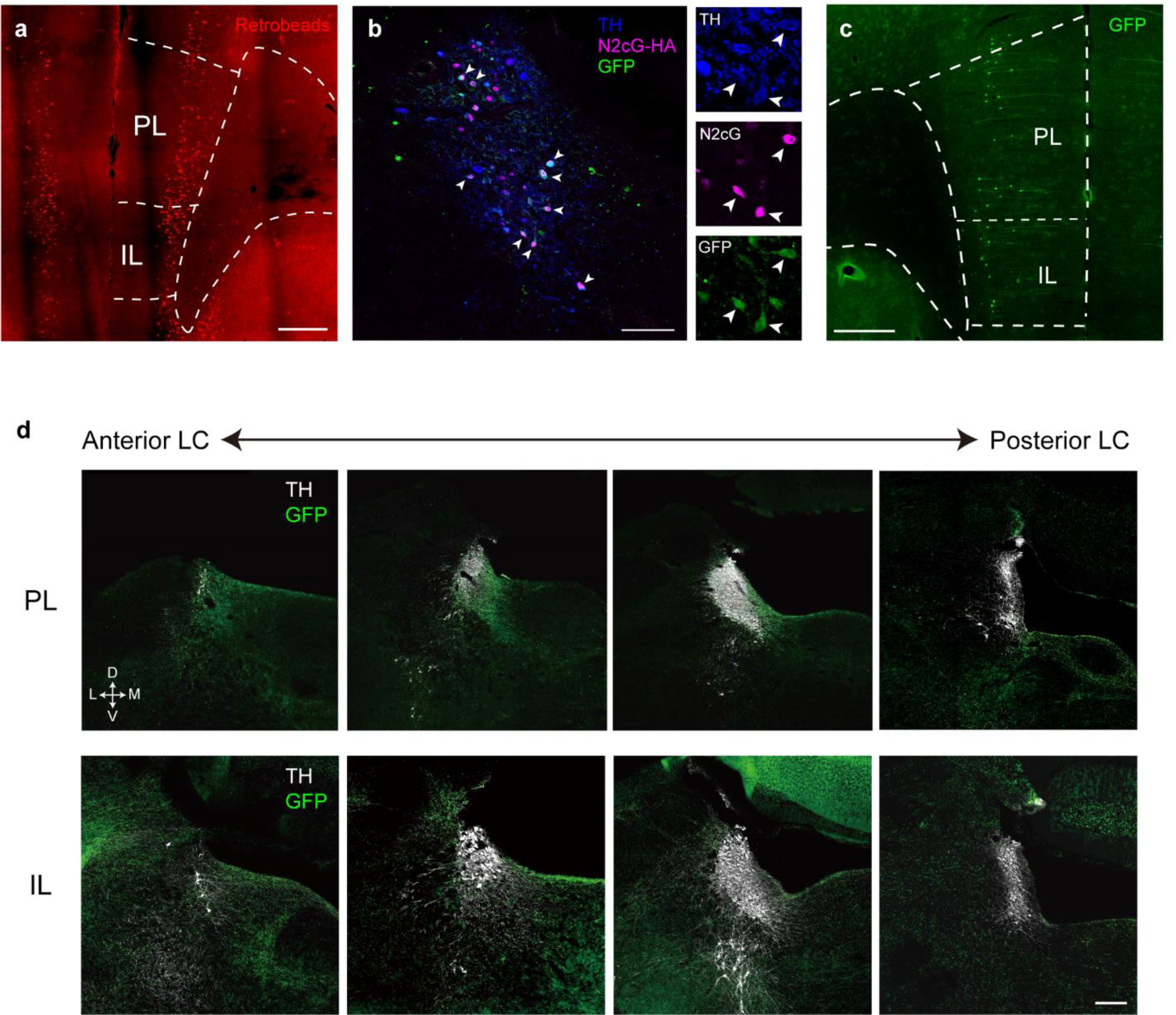
Anatomical connectivity from prefrontal subregions to LC-noradrenergic neurons. (***a***) Representative image for Retrobeads-labelled cells in PL and IL (Scale bar, 500 μm). (***b***) Representative image for rabies starter cells in LC. Arrowheads indicate starter cells expressing glycoprotein (N2cG) and rabies-GFP (Scale bar, 100 μm). Zoomed-in images on right show co-localization of TH, N2cG and GFP in representative starter cells. (***c***) Representative image for rabies input cells in PL and IL (Scale bar, 500 μm). (***d***) Spatial distributions of axon terminals from PL (top) or IL (bottom) through anterior-posterior axis of LC (Scale bar, 200 μm).

To further investigate the axonal projection pattern of PL and IL inputs to LC, we injected AAV carrying GFP into PL or IL of wild-type rats (*n*=3 for each). GFP-labeled axon terminals of PL and IL neurons were found in LC and pericoeruleus (peri-LC), which is adjacent to the LC and contains LC noradrenergic dendrites^25^ as well as distinct clusters of GABAergic neurons which regulate the activity of LC-noradrenergic neurons^15,26–28^ (***Fig. 1d***). We found that PL neurons predominantly project to the ventromedial region of peri-LC, while IL neurons project to dorsolateral as well as ventromedial peri-LC. These results reveal a distinct, topographically organized PL and IL anatomical innervation of LC, suggesting a mechanism through which PL and IL may differentially regulate LC noradrenergic activity.

### Bidirectional control of fear extinction learning by inputs from distinct mPFC subregions

Previous studies demonstrated that PL and IL have unique roles in the expression of emotional responding and fear extinction. Pharmacological and optogenetic studies revealed that IL activity is required for the formation of fear extinction memories^29–31^. By contrast, PL activation drives expression of aversive behavioral responses^32–34^ and electrophysiological studies showed that PL neural activity is negatively correlated with retrieval of fear extinction memories^35^, though its functional role in extinction is not clear. Based on this we hypothesized that IL inputs to LC would facilitate, while PL inputs to LC would oppose extinction of aversive emotional memories. To address this question, we performed optogenetic terminal inactivation during fear extinction learning. We injected AAV encoding the light-sensitive inhibitory archaerhodopsin with fluorescent proteins or fluorophore alone controls into either PL or IL and implanted fiber optics targeting the LC (***Fig. 2a-f***). Six-weeks after virus injection, rats underwent auditory fear conditioning, extinction learning, and retrieval sessions over three consecutive days. During extinction learning, laser illumination in LC was used to reduce terminal activity in the PL-to-LC and IL-to-LC pathways. Overlap groups received laser illumination during presentation of the conditioned stimulus (CS), while offset groups received laser for the same duration at pseudorandom timing during the inter-trial interval.

**Figure 2.**
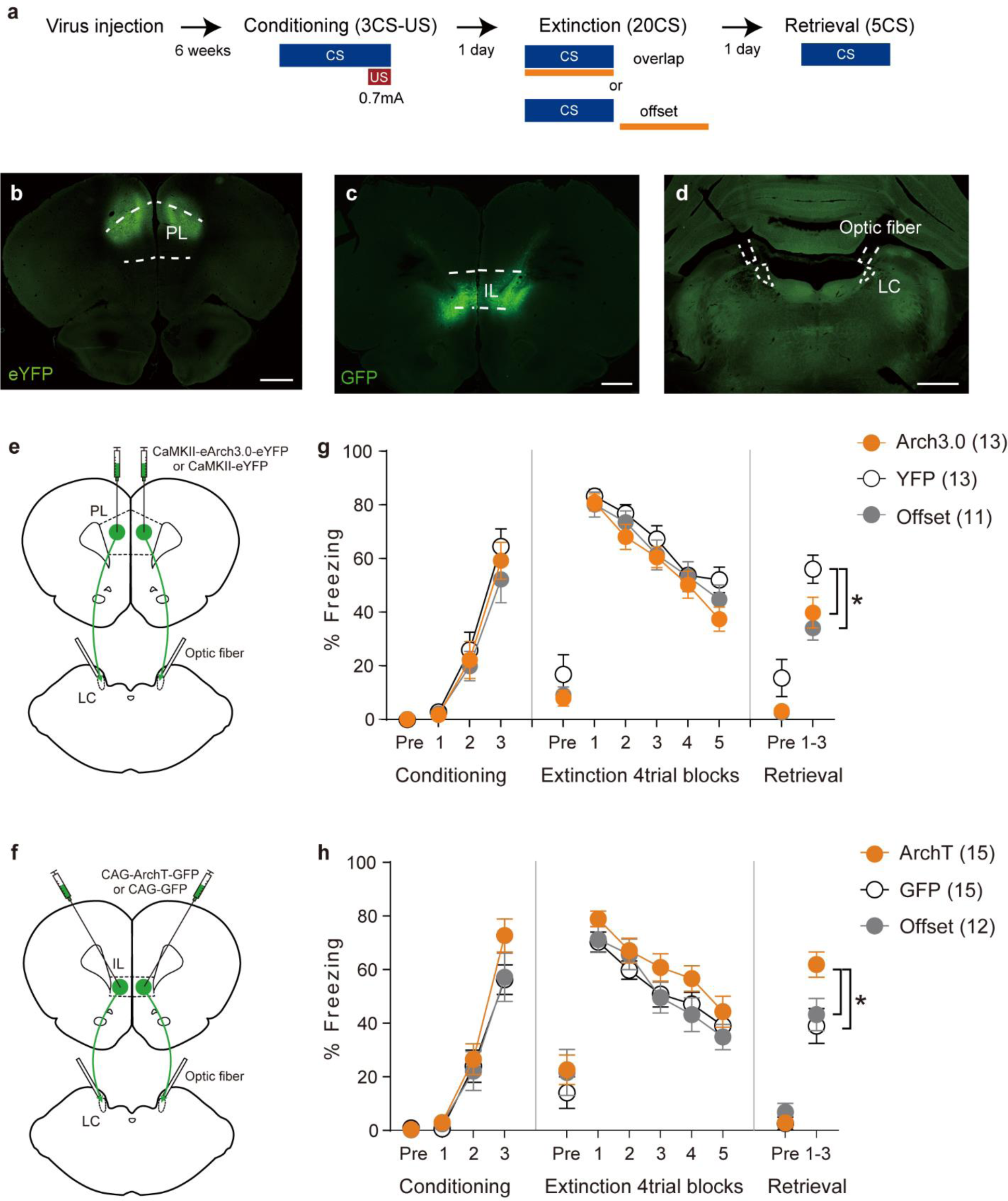
Prelimbic and infralimbic inputs to LC oppositely control formation of extinction memories. (***a***) Experimental schedule for optogenetic manipulation during fear extinction learning. (***b-c***) Representative images for virus expression in PL (***b***) and IL (***c***). (***d***) Representative image for the location of optical fiber targeting LC. (***e-f***) Schematic diagrams of virus injection and fiber implantation for manipulation of PL (***e***) and IL (***f***) inputs to LC. (***g***) Freezing during CS presentation and pre-CS baseline period (Pre) throughout fear conditioning, extinction learning and retrieval. PL-to-LC inactivation during CS or inter-trial-interval of extinction learning did not affect within-session freezing, but significantly enhanced extinction memory tested on the following day. (***h***) IL-to-LC inactivation during CS of extinction learning did not affect within-session freezing, but significantly impaired extinction memory tested on the following day. All scale bars, 1mm. All error bars indicate standard error of the mean (SEM) across animals. Numbers in brackets represent sample sizes. *p<0.05, one-way ANOVA & post-hoc Newman–Keuls test.

All groups acquired equivalent auditory fear memories during fear conditioning (***Fig. 2g-h***, two-way repeated measures ANOVA, PL-to-LC: group × trial interaction, F(4, 68) = 0.36, *p*=0.84, IL-to-LC: group × trial interaction, F(4, 78) = 0.94, *p*=0.44). Terminal inactivation of PL-to-LC and IL-to-LC inputs did not affect freezing reductions occurring during extinction learning (***Fig. 2g-h***, two-way repeated measures ANOVA, PL-to-LC: group × trial interaction, F(8, 136) = 0.75, *p*=0.65, IL-to-LC: group × trial interaction, F(8, 156) = 0.50, *p*=0.85), but produced opposing effects on the formation of long-term extinction memories which were tested on the following day. Inhibition of PL inputs to LC during the CS period or during the inter-trial-interval of extinction learning significantly enhanced long-term extinction memories relative to the fluorophore alone group (***Fig. 2g***, one-way ANOVA, F(2, 34) = 4.74, *p*=0.02). By contrast, inhibition of IL inputs to LC specifically during CS presentation of extinction learning significantly impaired formation of long-term extinction memories (***Fig. 2h***, one-way ANOVA, F(2, 39) = 4.67, *p*=0.02). These results demonstrate that IL-to-LC inputs are necessary to promote extinction memory formation, whereas activity in PL-to-LC inputs during the sensory CS period or during non-CS periods between trials oppose extinction.

### Distinct CS-evoked activity dynamics in LC-projecting PL and IL neurons during extinction learning

Based on the optogenetic experimental results showing that activity in the IL-to-LC pathway during the CS period facilitates extinction and previous findings in other systems showing ‘extinction neurons’ which have activity throughout or increasing during extinction training^9,36^, we hypothesized that IL-to-LC neurons would exhibit activity during periods of both high and low defensive responding during extinction while activity in PL-to-LC neurons would be specific to early extinction when defensive responding is high. To explore the activity dynamics of the PL-to-LC and IL-to-LC pathway neurons during fear extinction, we recorded calcium transients from retrogradely infected neurons using fiber photometry. We expressed a calcium indicator GCaMP6s in PL or IL neurons projecting to LC by injecting the combination of two retrograde viruses CAV2-Cre and AAV2retro-Cre into LC and AAV encoding cre-dependent GCaMP6s into PL or IL of wild-type rats and chronically implanted optical fibers in PL or IL (***Fig. 3a-c, f-g).*** For photometry recording, rats were habituated to fiber-optic patch cord tethering, exposed to auditory CSs during a pre-conditioning habituation session and then underwent fear conditioning, extinction learning and memory retrieval sessions (***Fig. 3a, Supplementary Fig. 1***).

**Figure 3.**
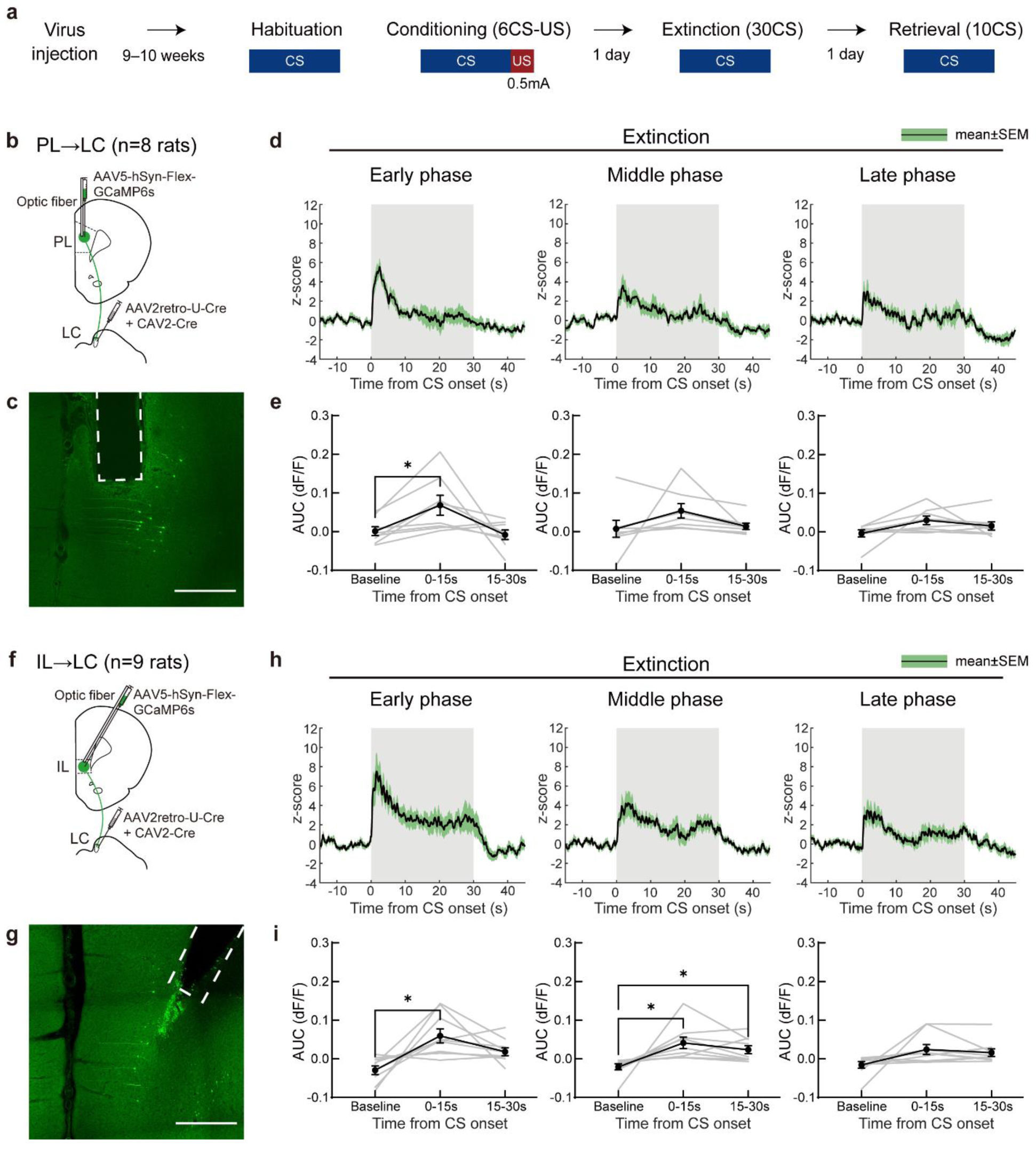
Distinct calcium dynamics in prelimbic and infralimbic inputs to LC during fear extinction learning. (***a***) Experimental schedule for fiber photometry recording. (***b***) Schematic of virus injections and fiber implantation for PL-to-LC inputs. (***c***) Representative image for GCaMP expression in PL neurons. Dotted lines indicate tracks of optic fibers. (***d***) CS-triggered calcium responses in PL-to-LC inputs during early (trial 1-10), middle (trial 11-20), and late (trial 21-30) phases of extinction learning. Black lines show average and green shades show SEM across animals. Shaded areas in grey indicate CS-presentation time period. (***e***) Area under curve (AUC) of PL-to-LC dF/F in pre-CS baseline period (BL, -15 to 0 s) and CS presentation periods (0-15 s and 15-30 s). (***f***) Schematic of virus injections and fiber implantation for IL-to-LC inputs. (***g***) Representative image for GCaMP expression in IL neurons. (***h***) CS-triggered calcium responses in IL-to-LC inputs during early, middle, and late phases of extinction learning. (***i***) AUC of IL-to-LC dF/F in pre-CS baseline period (BL, -15 to 0 s) and CS presentation periods (0-15 s and 15-30 s). All scale bars, 500 μm. All error bars indicate SEM. *p<0.05, **p<0.01 (one-way repeated ANOVA and Dunnett’s multiple comparison test). All error bars indicate SEM across animals.

Examining sensory evoked CS-responses, we found that PL-to-LC and IL-to-LC input neurons had distinct calcium dynamics during fear extinction learning (***Fig. 3d,h***). PL-to-LC neurons responded phasically at CS onset in the beginning of extinction learning, and this CS-evoked phasic activation diminished in later trials (***Fig. 3d-e***, one-way repeated measures ANOVA, early phase: F(1.30, 9.10) = 5.02, P=0.04, middle phase: F(1.11, 7.78) = 2.54, *p*=0.15, late phase: F(1.71, 12.0) = 2.16, *p*=0.16). By contrast, IL-to-LC neurons showed more sustained activation during CS presentation than PL-to-LC neurons and this activation was more prolonged, occurring in both early and later trials of extinction training (***Fig. 3h-i***, one-way repeated measures ANOVA, early phase: F(1.66, 13.3) = 7.75, *p*=0.008, middle phase: F(1.39, 11.1) = 6.56, *p*=0.02, late phase: F(1.32, 10.6) = 3.20, *p*=0.09). Together, the differential activity dynamics in PL-to-LC and IL-to-LC inputs suggest distinct processes in regulating fear extinction.

### PL-to-LC neuronal activity is related to internally generated aversive states during extinction learning

Electrophysiological studies reported that PL neurons encode freezing information including internally generated, spontaneous freezing outside of the CS period^37^. Furthermore, our optogenetic results demonstrated that inactivation of PL-to-LC inputs during CS and inter-trial-interval periods enhanced extinction memory formation (***Fig. 2g***). Based on these findings, we hypothesized that activation of PL-to-LC inputs may be driven by internal aversive states occurring during freezing episodes. To test this hypothesis, we analyzed activity in LC-projecting PL and IL neurons in relation to freezing behaviors during the CS period and the inter-trial-interval. We found that PL-to-LC neurons exhibited significant calcium increases at freezing onset compared to the baseline period (***Fig. 4a-b***, one-way repeated measures ANOVA, F(2.32, 473.6) = 7.94, *p*=0.0002). Calcium activity significantly increased prior to the onset of freezing during non-CS periods (***Fig. 4e-f***, one-way repeated measures ANOVA, F(2.41, 223.9) = 5.66, *p*=0.002) and were high prior to freezing onset during CS periods (***Fig. 4g-h***), likely reflecting combined CS and freezing related activity. Since the activation of PL-to-LC inputs began prior to freezing onset (***Fig. 4b,f***), it is not likely to reflect feedback from freezing behavior but rather encode an internal freezing command or aversive state. By contrast, IL-to-LC inputs showed no activation at freezing onset (one-way repeated measures ANOVA, all freezing episodes: ***Fig. 4c-d***, F(2.41, 584.5) = 1.65, *p*=0.19 outside CS: ***Fig. 4i-j***, F(1.69, 199.8) = 1.09, *p*=0.33, during CS: ***Fig. 4k-l***, F(3.09, 382.8) = 0.94, *p*=0.42). Together, these results demonstrate that activation of PL-to-LC, but not IL-to-LC, projection neurons is time-locked to behavioral freezing. This freezing-related activation of PL-to-LC inputs may be important for state-dependent control of fear extinction to constrain extinction memory formation in highly aversive states.

**Figure 4.**
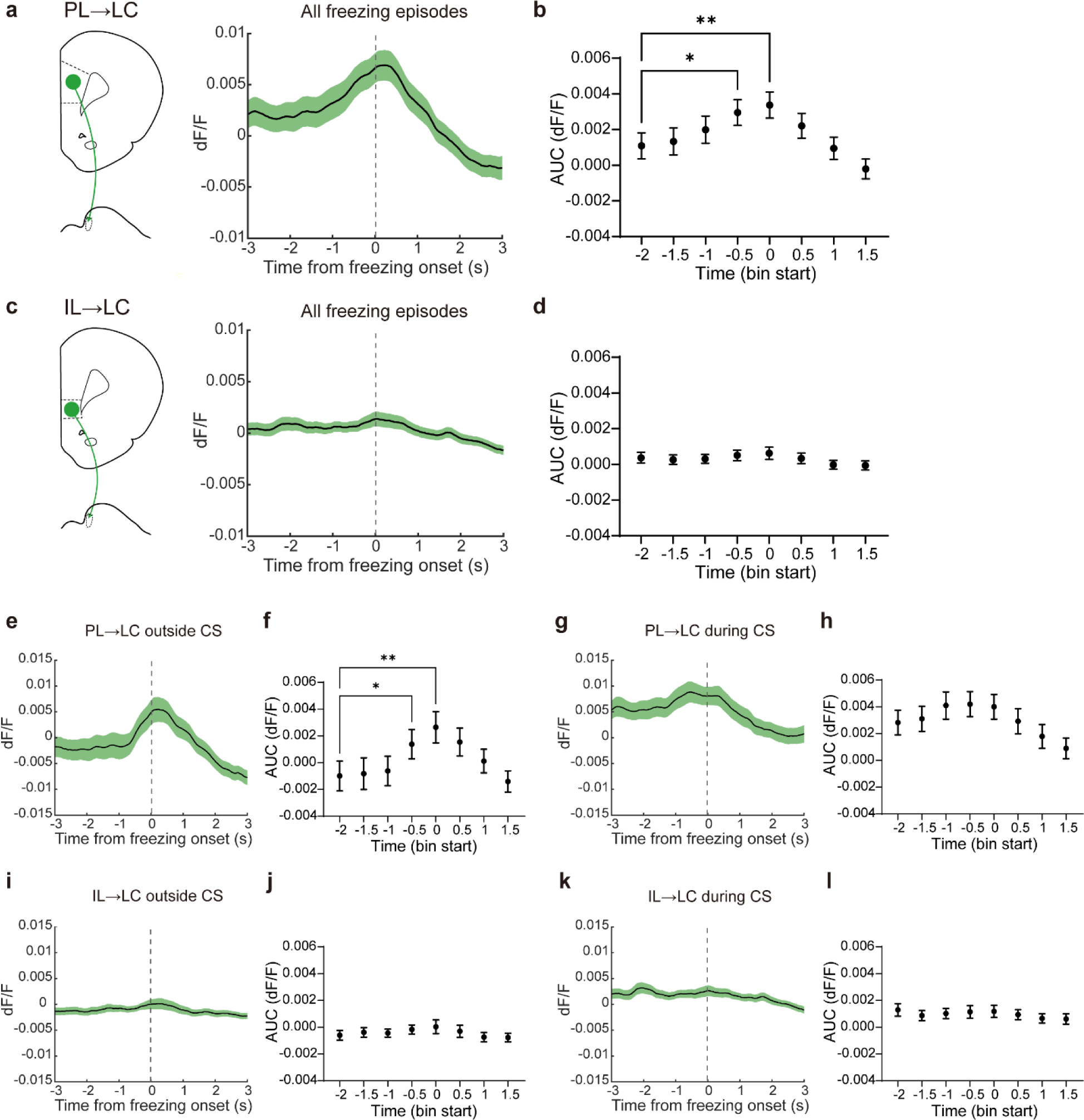
Freezing-locked calcium increases in PL-to-LC but not IL-to-LC pathway neurons. (***a,c***) Freezing-triggered calcium responses in PL-to-LC (***a***) and IL-to-LC (***c***) inputs. Black lines and green shades show average and SEM, respectively. (***b,d***) Comparison of AUCs of freezing-triggered dF/F in 0.5-s time bin with pre-freezing baseline period (−2 to −1.5 s). (***e-f***) Freezing-triggered calcium responses in PL-to-LC inputs outside CS. (***g-h***) Freezing-triggered calcium responses in PL-to-LC inputs during CS. (***i-j***) Freezing-triggered calcium responses in IL-to-LC inputs outside CS. (***k-l***) Freezing-triggered calcium responses in IL-to-LC inputs during CS. All error bars indicate SEM. *p<0.05, ** p<0.01 (one-way repeated ANOVA and Dunnett’s multiple comparison test).

### Discussion

Here we show that PL and IL prefrontal subregions opposingly regulate extinction of aversive emotional memories through distinct projections to the LC-noradrenaline system. Anatomical tracing experiments demonstrated that the LC-noradrenergic neurons receive synaptic inputs from PL and IL and that these subregions have unique axonal projection patterns in peri-LC. Optogenetic manipulation experiments revealed that PL-to-LC and IL-to-LC inputs function to suppress or promote formation of long-term extinction memories, respectively. Fiber photometry calcium imaging experiments identified distinct activity dynamics in these inputs: PL-to-LC projection neurons responded to sensory cues during high aversive states early in extinction and to internally generated freezing behavior, while IL-to-LC projection neurons showed sustained cue-evoked activation during early and middle phases of extinction learning. These findings suggest that PL and IL provide state-dependent, top-down control of emotional processing through innervation of the LC-noradrenaline system.

While noradrenergic modulation of mPFC has been examined extensively, the role of prefrontal influences on the noradrenaline system is not well understood. Noradrenaline in mPFC is known to modulate emotional expression and extinction as well as cognitive processes such as behavioral flexibility and decision making^3,9,38–44^. PL and IL themselves contribute to distinct emotional functions, with PL being important for expression of various defensive responses while IL facilitates extinction of aversive memories^34,45–47^. Our findings here extend this idea by showing that aversive emotional information is sent from PL to the LC to restrain extinction memory formation while IL projections to LC convey pro-extinction information.

How PL and IL information is integrated in LC to regulate the level of noradrenaline release throughout the brain and in individual brain regions is an important open question. Prior work has shown that unique subpopulations of LC-noradrenaline neurons have specific connectivity with efferent target structures, serve distinct behavioral functions and exhibit activity profiles distinct from the larger LC-noradrenaline neural pool^9,38,42,48–51^. However, many LC neurons project widely throughout the brain and LC-noradrenaline neurons respond uniformly in many situations^9,38,50,52^. It is still unclear whether PL and IL inputs affect global LC activity or exert their influences through specific noradrenergic subpopulations. Previous studies demonstrated that projection-defined subpopulations in LC-noradrenergic neurons are differentially involved in emotional processing^9,10,42,48^. Notably, different populations of LC-noradrenaline neurons project to the amygdala or mPFC and oppose or facilitate, respectively, fear extinction^9^. Though amygdala-projecting and mPFC-projecting neurons in LC receive similar inputs from PL and IL^53^, it does not exclude the possibility that PL and IL differentially influence neural activity in specific populations of LC-noradrenaline neurons via multi-synaptic connections through peri-LC GABAergic neurons or more subtle differences in synaptic strength in PL and IL inputs to these LC-noradrenergic neuronal subpopulations. Indeed, LC-noradrenergic activity can be inhibited through OFC-to-peri-LC-to-LC disynaptic connections mediating arousal control^15^. Furthermore, a recent study reported that distinct clusters of peri-LC GABAergic neurons differentially respond to emotionally arousing stimuli^54^. Further investigations on connectivity and activity recording from LC/peri-LC local circuits are required to determine more detailed mechanisms through which mPFC exerts top-down control of LC-noradrenergic activity in fear extinction.

### Methods

#### Animals

All experimental procedures were approved by the Animal Care and Use Committees of the RIKEN Center for Brain Science. Male adult Long-Evans (8–12 weeks old at the time of surgery) or TH-Cre (9 weeks old at the time of surgery) rats were used for all experiments. Animals were singly housed on a 12-h light/dark cycle, and food and water were provided ad libitum. All behavioral experiments were performed during the light cycle.

#### Viruses

Adeno-associated viruses (AAV, serotype 5) containing CAG-ArchT-GFP, CAG-GFP, CaMKII-eArch3.0-eYFP, and CaMKII-eYFP were obtained from the University of North Carolina Vector Core. Canine adenovirus (CAV2)-Cre was provided by the Montpellier vector core. Plasmid of pAAV-CAG-flex-HA-N2cG was obtained from Addgene (#73477) and packaged into AAV9/2 in our laboratory. AAV9/2-CBA-flex-TVA was produced and packaged in our laboratory. Plasmid for EnvA-coated CVS-strain glycoprotein-deleted rabies virus fused with EGFP (EnvA-CVS-N2cΔG-EGFP) ^24^ was provided by Dr. Thomas Jessell (Addgene #73461) and packaged in our laboratory using the Neuro2A cell lines (mouse neuroblasts) provided by Dr. Andrew Murray. Retrograde AAV (setotype 2)-U-Cre (AAV2retro-U-Cre) was produced and packaged in our laboratory. AAV5-hSyn-flex-GCaMP6s was obtained from Addgene (#100845).

#### Surgery

Rats were anesthetized with isoflurane (5% for induction, 1.5–2.5% for maintenance) and placed in a stereotaxic frame (Kopf Instruments). For retrograde tracing, 0.2 µL of red Retrobeads (Lumafluor Inc.) was unilaterally injected into LC (30°, AP Interaural−1.1, ML ±4.55, DV −5.5). For transsynaptic rabies tracing, 1.0 µL of helper AAVs carrying N2cG and TVA (mixed 1:1) were unilaterally injected into the LC (30°, AP Interaural−1.1, ML ±4.55, DV −5.5) and 1.5 µL of rabies EnvA-CVS-N2cΔG-GFP was injected into the LC (same coordinates) three weeks after the helper injection. For anterograde tracing and optogenetic experiments, 0.5 µL of AAV was bilaterally injected into PL (0°, AP 2.9, ML ±0.55, DV −3.1) or IL (30°, AP 2.75, ML ±2.95, DV −4.2). For optogenetic experiments, fiber-optic cannulas (Doric Lenses, 200 µm, NA 0.37) were bilaterally implanted in the LC (30°, AP Interaural−1.1, ML ±4.55, DV −5.2). For fiber photometry experiments, 0.4 µL of AAV2retro-U-Cre and CAV2-Cre (mixed 1:1) were unilaterally injected into the LC (30°, AP Interaural−1.1, ML 4.55, DV −5.5) and 0.5 µL of AAV5-hSyn-flex-GCaMP6s was injected into PL (0°, AP 2.9, ML 0.55, DV −3.1) or IL (30°, AP 2.75, ML 2.95, DV −4.2) of the same hemisphere as LC. Fiber-optic cannula (Doric Lenses, 400 µm, NA 0.66) was implanted in PL or IL and secured using stainless steel screws, UV-sensitive glue, superbond and dental cement.

#### Optogenetic manipulation during fear extinction

Rats were handled for two days prior to fear conditioning. Rats were placed in a sound-isolated chamber (Med Associates, Inc.) and presented with stimuli. On day 1, rats were conditioned with three presentations of an auditory conditioned stimulus (CS; 5 kHz pip, 20 s) which co-terminated with an aversive unconditioned stimulus (US; electrical footshock, 0.7 mA, 1s) in a conditioning context. On day 2, the rats were exposed to 20 CS presentations without footshock in a novel context for extinction training. On day 3, the rats were presented with five CSs in the extinction context to test long-term extinction memory.

Behavioral freezing responses during CS presentation were scored manually by an experimenter blind to the treatment conditions. Context freezing was also scored for 20 seconds before the first CS presentation on each day.

For optogenetic inhibition, 589-nm orange laser was delivered through the implanted fiber optics during extinction training. Arch and fluorophore (GFP or eYFP) control groups received laser during CS presentation, which was turned on 400 ms before CS onset and turned off 50 ms after CS offset. Offset control group received laser for the same duration at pseudorandom timings after every CS presentation.

#### Fiber photometry recording

Photometry recordings were conducted 9-10 weeks after GCaMP injection. Rats were handled for two days and habituated to being tethered to the patch cord prior to recordings. Animals were placed in a behavioral chamber (Med Associates, Inc.) and delivered an auditory conditioned stimulus (CS; 5 kHz pip, 30 s) and an unconditioned stimulus (US; electrical footshock, 0.5 mA, 2s).

On day 1, in a conditioning context, rats were first exposed to four or six CSs for habituation, and then fear conditioned with six CS-US parings. On day 2, rats were exposed to 30 CS trials without footshock in a novel context for extinction training. On day 3, the rats were presented with 10 CSs in the extinction context to test long-term extinction memory.

Calcium transients were recorded with a Doric fiber photometry system (Doric Lenses) measuring 405-nm isosbestic and 470-nm GCaMP-dependent signals. Signals were sampled at 12 kHz. Video frames were triggered at 30 Hz by Doric Neuroscience Studio to synchronize with photometry recordings.

#### Photometry data analysis

All photometry data were analyzed using custom MATLAB scripts. Signals were first downsampled to 1 kHz. To calculate ΔF/F, the isosbestic signal (405 nm) was aligned to the GCaMP-dependent signal (470 nm) using a least-squares linear fit to adjust distinct damping curves of the two signals. The fitted isosbestic signal was then subtracted from the GCaMP-dependent signal. ΔF/F was smoothed using a moving average with 200-ms time window for the later analysis.

For CS-evoked activity plots, z-score was calculated using pre-CS 15-s baseline period. AUC of ΔF/F was calculated during baseline (-15 to 0 s from CS onset), first half of CS (0 to 15 s), and the second half of CS (15 to 30 s) periods.

For freezing-locked activity analysis, freezing episodes were manually labeled at single video frame resolution using BENTO, a MATLAB-based graphical user interface^55^. Only freezing episodes that lasted for at least 1 s in the middle phase of extinction learning (from trial 11 to 20) were included in the analysis. Freezing episodes with freezing in the preceding 0.5-s period prior to their onset or unclear onset (e.g. due to visual occlusion) were excluded from the analysis. AUC of ΔF/F was calculated in 0.5-s time bins.

#### Histology

To evaluate virus expression and fiber location, rats were intraperitoneally overdosed with 3.0 mL of 25% chloral hydrate and transcardially perfused with 4% paraformaldehyde in PBS. After overnight post-fixation, brains were sliced into 40 or 50-µm coronal sections using a cryostat.

#### Immunohistochemistry

Brain sections were blocked in 2% donkey serum in PBST for 30 minutes and incubated in primary antibodies overnight at 4 °C in the dark. Sections were then rinsed in PBS and incubated in secondary antibodies for 1 hour. After rinsing with PBS, sections were mounted on slides and coverslipped with Fluoromount Plus (Diagnostic BioSystems Inc.).

For anterograde tracing, primary mouse anti-GFP (Thermo Fisher Scientific, A-11120, 1:2000) and rabbit anti-tyrosine hydroxylase (TH) (Millipore, AB152, 1:2000) antibodies were used with secondary donkey Alexa Fluor 488 anti-mouse and Alexa Fluor 594 anti-rabbit antibodies (Thermo Fisher Scientific, A-21202 and 21207, 1:1000).

For monosynaptic rabies tracing experiments, primary mouse anti-GFP (same as above), rabbit anti-HA (Cell Signaling technology, #3724, 1:5000), and sheep anti-TH (Millipore, AB1542, 1:2000) were used with secondary donkey Alexa Fluor 488 anti-mouse, Alexa Fluor 594 anti-rabbit (same as above) and Alexa Fluor 647 anti-sheep (Thermo Fisher Scientific, A-21448, 1:1000) antibodies.

#### Statistical analysis

Statistical analyses were performed using GraphPad Prism (GraphPad Software, Inc.). Two-way repeated measures ANOVA or one-way ANOVA were used to detect differences in freezing between experimental groups. Significant main effects of one-way ANOVA were followed by Newman-Keuls post-hoc tests.

One-way repeated measures ANOVA with Geisser-Greenhouse’s correction was used to detect differences in calcium activities between time bins. Significant main effects of repeated measures ANOVA were followed by Dunnet post-hoc tests. The significance level was set at P < 0.05 for all results.

## Data and code availability

The data and code that support the findings of this study are available from the corresponding author upon reasonable request.

## Acknowledgements

We thank Yuri Sugiyama, Reiko Yoshida, and Nisha Jose for technical assistance. We thank Dr. Andrew Murray for gifting the Neuro2A cell lines used for packaging CVS-strain rabies virus. This work was supported by funding from the RIKEN Center for Brain Science, RIKEN Junior Research Associate Program and KAKENHI 19H05234.

## Author contributions

M.W., A.U., and J.P.J. conceived the project and designed the experiments. M.W. and A.U. carried out the optogenetics experiments. M.W. carried out the anatomical tracing and fiber photometry experiments and analyzed all data. M.W. and J.P.J. wrote the manuscript. All authors read and approved the final manuscript.

---

**Supplementary Figure 1.**
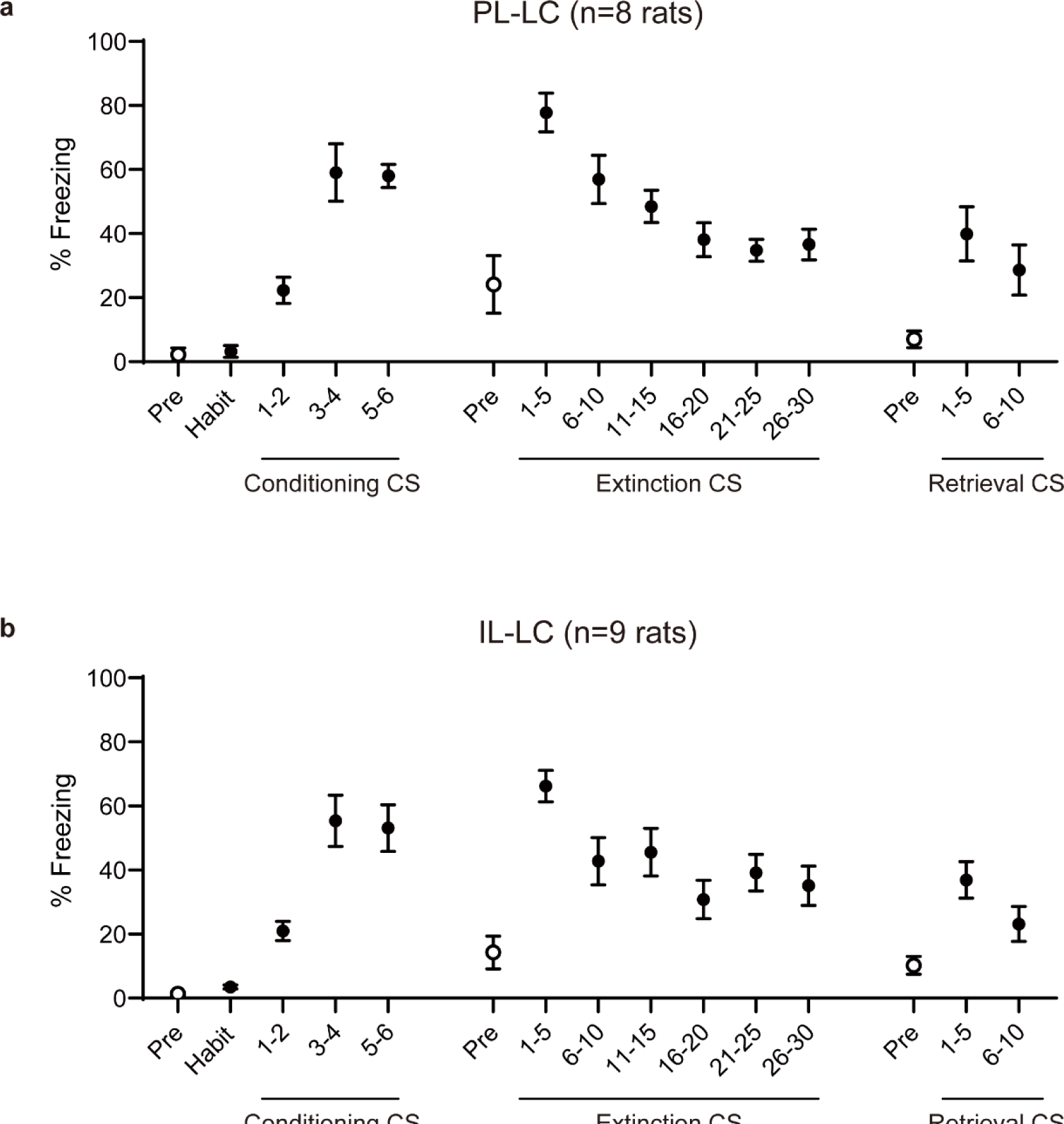
Freezing during fiber photometry recording. Freezing of PL-to-LC rats (***a***) and IL-to-LC rats (***b***) during CS presentation and pre-CS baseline period (Pre) throughout habituation, fear conditioning, extinction learning, and retrieval.

**Supplementary Figure 2.**
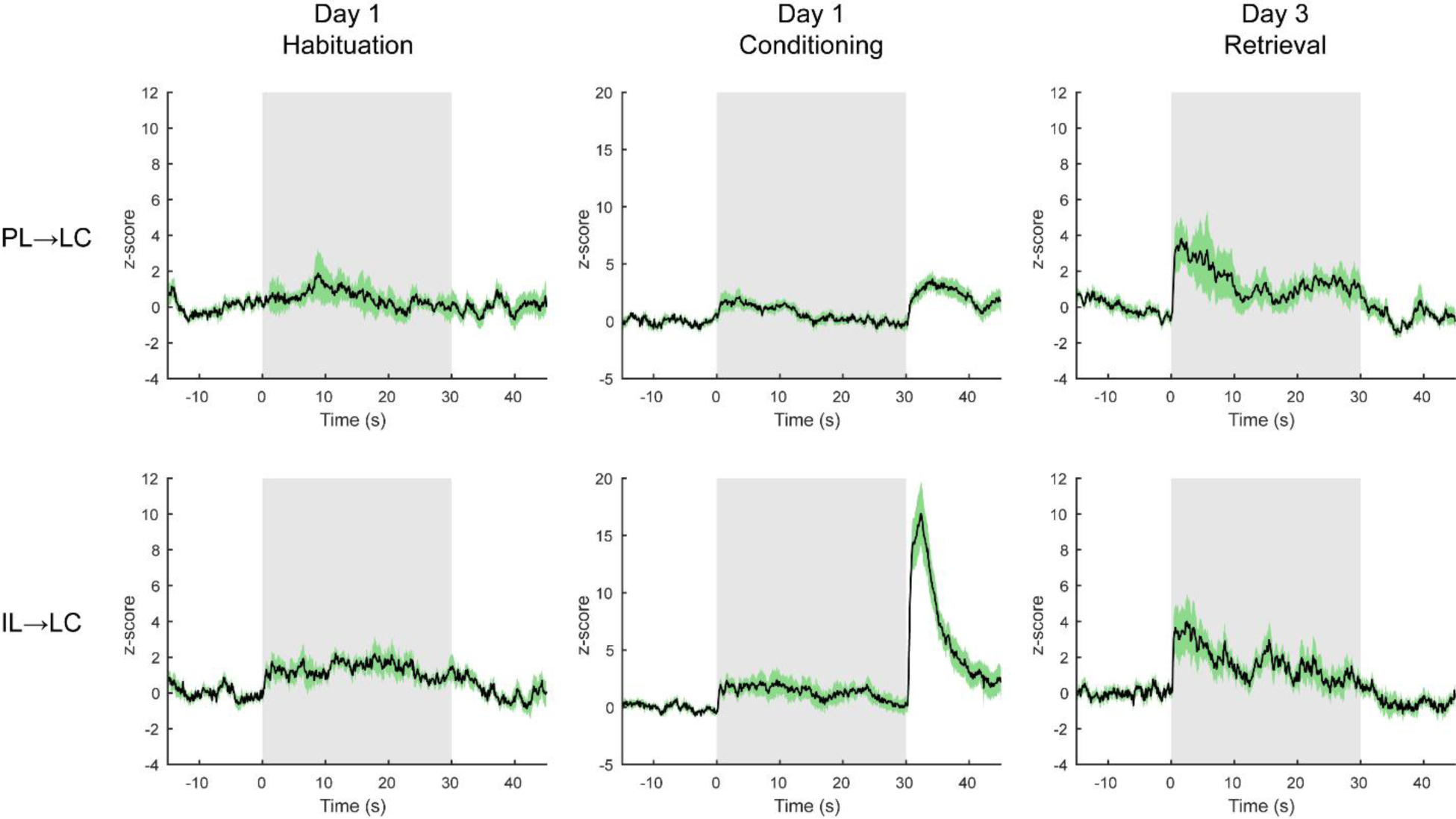
CS-evoked calcium responses during habituation, fear conditioning, and retrieval. CS-triggered calcium responses in PL-to-LC (top) and IL-to-LC (bottom) inputs. Black lines show average and green shades show SEM across animals. Shaded areas in grey indicate CS-presentation time period.

## References

1. Bush, D. E. A., Caparosa, E. M., Gekker, A. & Ledoux, J. Beta-adrenergic receptors in the lateral nucleus of the amygdala contribute to the acquisition but not the consolidation of auditory fear conditioning. Front Behav Neurosci 4, 154 (2010).

2. Schiff, H. C. et al. β-Adrenergic Receptors Regulate the Acquisition and Consolidation Phases of Aversive Memory Formation Through Distinct, Temporally Regulated Signaling Pathways. Neuropsychopharmacology 42, 895–903 (2017).

3. Mueller, D., Porter, J. T. & Quirk, G. J. Noradrenergic Signaling in Infralimbic Cortex Increases Cell Excitability and Strengthens Memory for Fear Extinction. The Journal of Neuroscience 28, 369– 375 (2008).

4. Likhtik, E. & Johansen, J. P. Neuromodulation in circuits of aversive emotional learning. Nat Neurosci 22, 1586–1597 (2019).

5. Poe, G. R. et al. Locus coeruleus: a new look at the blue spot. Nat Rev Neurosci 21, 644–659 (2020).

6. Giustino, T. F. & Maren, S. Noradrenergic Modulation of Fear Conditioning and Extinction. Front Behav Neurosci 12, 43 (2018).

7. Yeh, L., Watanabe, M., Sulkes-cuevas, J. & Johansen, J. P. Dysregulation of aversive signaling pathways: a novel circuit endophenotype for pain and anxiety disorders. Curr Opin Neurobiol 48, 37–44 (2018).

8. Southwick, S. M. et al. Role of norepinephrine in the pathophysiology and treatment of posttraumatic stress disorder. Biol Psychiatry 46, 1192–1204 (1999).

9. Uematsu, A. et al. Modular organization of the brainstem noradrenaline system coordinates opposing learning states. Nat Neurosci 20, 1602–1611 (2017).

10. McCall, J. G. et al. Locus coeruleus to basolateral amygdala noradrenergic projections promote anxiety-like behavior. Elife 6, e18247 (2017).

11. Haubrich, J., Bernabo, M. & Nader, K. Noradrenergic projections from the locus coeruleus to the amygdala constrain fear memory reconsolidation. Elife 9, e57010 (2020).

12. Fitzgerald, P. J., Giustino, T. F., Seemann, J. R. & Maren, S. Noradrenergic blockade stabilizes prefrontal activity and enables fear extinction under stress. Proc Natl Acad Sci U S A 112, E3729– E3737 (2015).

13. Soya, S. et al. Orexin modulates behavioral fear expression through the locus coeruleus. Nat Commun 8, 1606 (2017).

14. Luppi, P. H., Aston-Jones, G., Akaoka, H., Chouvet, G. & Jouvet, M. Afferent projections to the rat locus coeruleus demonstrated by retrograde and anterograde tracing with cholera-toxin B subunit and Phaseolus vulgaris leucoagglutinin. Neuroscience 65, 119–160 (1995).

15. Breton-Provencher, V. & Sur, M. Active control of arousal by a locus coeruleus GABAergic circuit. Nat Neurosci 22, 218–228 (2019).

16. Lu, Y., Simpson, K. L., Weaver, K. J. & Lin, R. C. S. Differential Distribution Patterns From Medial Prefrontal Cortex and Dorsal Raphe to the Locus Coeruleus in Rats. Anat Rec 295, 1192–1201 (2012).

17. Aston-Jones, G. & Cohen, J. D. An Integrative Theory of Locus Coeruleus-Norepinephrine Function: Adaptive Gain and Optimal Performance. Annu Rev Neurosci 28, 403–450 (2005).

18. Jodo, E., Chiang, C. & Aston-Jones, G. Potent excitatory influence of prefrontal cortex activity on noradrenergic locus coeruleus neurons. Neuroscience 83, 63–79 (1998).

19. Sara, S. J. & Hervé-Minvielle, A. Inhibitory influence of frontal cortex on locus coeruleus neurons. Proc Natl Acad Sci U S A 92, 6032–6036 (1995).

20. Barcomb, K., Olah, S. S., Kennedy, M. J. & Ford, C. P. Properties and modulation of excitatory inputs to the locus coeruleus. J Physiol 600, 4897–4916 (2022).

21. Totah, N. K., Logothetis, N. K. & Eschenko, O. Synchronous spiking associated with prefrontal high γ oscillations evokes a 5-Hz rhythmic modulation of spiking in locus coeruleus. J Neurophysiol 125, 1191–1201 (2021).

22. Cardenas, A., Papadogiannis, A. & Dimitrov, E. The role of medial prefrontal cortex projections to locus ceruleus in mediating the sex differences in behavior in mice with inflammatory pain. FASEB Journal 35, e21747 (2021).

23. Witten, I. B. et al. Recombinase-driver rat lines: Tools, techniques, and optogenetic application to dopamine-mediated reinforcement. Neuron 72, 721–733 (2011).

24. Reardon, T. R. et al. Rabies Virus CVS-N2cΔG Strain Enhances Retrograde Synaptic Transfer and Neuronal Viability. Neuron 89, 711–724 (2016).

25. Shipley, M. T., Fu, L., Ennis, M., Liu, W. L. & Aston-Jones, G. Dendrites of locus coeruleus neurons extend preferentially into two pericoerulear zones. Journal of Comparative Neurology 365, 56–68 (1996).

26. Aston-Jones, G., Zhu, Y. & Card, J. P. Numerous GABAergic Afferents to Locus Ceruleus in the Pericerulear Dendritic Zone: Possible Interneuronal Pool. Journal of Neuroscience 24, 2313–2321 (2004).

27. Jin, X. et al. Identification of a group of GABAergic neurons in the dorsomedial area of the locus coeruleus. PLoS One 11, e0146470 (2016).

28. Kuo, C.-C. et al. Inhibitory interneurons regulate phasic activity of noradrenergic neurons in the mouse locus coeruleus and functional implications. Journal of Physiology 598, 4003–4029 (2020).

29. Do-Monte, F. H., Manzano-Nieves, G., Quinones-Laracuente, K., Ramos-Medina, L. & Quirk, G. J. Revisiting the Role of Infralimbic Cortex in Fear Extinction with Optogenetics. Journal of Neuroscience 35, 3607–3615 (2015).

30. Bukalo, O. et al. Effects of optogenetic photoexcitation of infralimbic cortex inputs to the basolateral amygdala on conditioned fear and extinction. Behavioural Brain Research 396, 112913 (2021).

31. Quirk, G. J., Garcia, R. & González-Lima, F. Prefrontal Mechanisms in Extinction of Conditioned Fear. Biol Psychiatry 60, 337–343 (2006).

32. Courtin, J. et al. Prefrontal parvalbumin interneurons shape neuronal activity to drive fear expression. Nature 505, 92–96 (2014).

33. Do-Monte, F. H., Quiñones-Laracuente, K. & Quirk, G. J. A temporal shift in the circuits mediating retrieval of fear memory. Nature 519, 460–463 (2015).

34. Corcoran, K. A. & Quirk, G. J. Activity in prelimbic cortex is necessary for the expression of learned, but not innate, fears. Journal of Neuroscience 27, 840–844 (2007).

35. Burgos-Robles, A., Vidal-Gonzalez, I. & Quirk, G. J. Sustained conditioned responses in prelimbic prefrontal neurons are correlated with fear expression and extinction failure. Journal of Neuroscience 29, 8474–8482 (2009).

36. Herry, C. et al. Switching on and off fear by distinct neuronal circuits. Nature 454, 600–606 (2008).

37. Dejean, C. et al. Prefrontal neuronal assemblies temporally control fear behaviour. Nature 535, 420–424 (2016).

38. Breton-Provencher, V., Drummond, G. T., Feng, J., Li, Y. & Sur, M. Spatiotemporal dynamics of noradrenaline during learned behaviour. Nature 606, 732–738 (2022).

39. Giustino, T. F., Fitzgerald, P. J., Ressler, R. L. & Maren, S. Locus coeruleus toggles reciprocal prefrontal firing to reinstate fear. Proc Natl Acad Sci U S A 116, 8570–8575 (2019).

40. Do-Monte, F. H. M., Allensworth, M. & Carobrez, A. P. Impairment of contextual conditioned fear extinction after microinjection of alpha-1-adrenergic blocker prazosin into the medial prefrontal cortex. Behavioural Brain Research 211, 89–95 (2010).

41. Bari, A. et al. Differential attentional control mechanisms by two distinct noradrenergic coeruleo-frontal cortical pathways. Proc Natl Acad Sci U S A 117, 29080–29089 (2020).

42. Borodovitsyna, O., Duffy, B. C., Pickering, A. E. & Chandler, D. J. Anatomically and functionally distinct locus coeruleus efferents mediate opposing effects on anxiety-like behavior. Neurobiol Stress 13, 100284 (2020).

43. Arnsten, A. F. T. Stress signalling pathways that impair prefrontal cortex structure and function. Nat Rev Neurosci 10, 410–422 (2009).

44. Cope, Z. A., Vazey, E. M., Floresco, S. B. & Aston Jones, G. S. DREADD-mediated modulation of locus coeruleus inputs to mPFC improves strategy set-shifting. Neurobiol Learn Mem 161, 1–11 (2019).

45. Jercog, D. et al. Dynamical prefrontal population coding during defensive behaviours. Nature 595, 690–694 (2021).

46. Maren, S. & Holmes, A. Stress and Fear Extinction. Neuropsychopharmacology 41, 58–79 (2016).

47. Sotres-Bayon, F. & Quirk, G. J. Prefrontal control of fear: more than just extinction. Curr Opin Neurobiol 20, 231–235 (2010).

48. Hirschberg, S., Li, Y., Randall, A., Kremer, E. J. & Pickering, A. E. Functional dichotomy in spinal-vs prefrontal-projecting locus coeruleus modules splits descending noradrenergic analgesia from ascending aversion and anxiety in rats. Elife 6, e29808 (2017).

49. Totah, N. K., Neves, R. M., Panzeri, S., Logothetis, N. K. & Eschenko, O. The Locus Coeruleus Is a Complex and Differentiated Neuromodulatory System. Neuron 99, 1055–1068.e6 (2018).

50. Kebschull, J. M. et al. High-Throughput Mapping of Single-Neuron Projections by Sequencing of Barcoded RNA. Neuron 91, 975–987 (2016).

51. Chandler, D. J., Gao, W.-J. & Waterhouse, B. D. Heterogeneous organization of the locus coeruleus projections to prefrontal and motor cortices. Proc Natl Acad Sci U S A 111, 6816–21 (2014).

52. Schwarz, L. A. et al. Viral-genetic tracing of the input-output organization of a central noradrenaline circuit. Nature 524, 88–92 (2015).

53. Sulkes Cuevas, J. N., Watanabe, M., Uematsu, A. & Johansen, J. P. Whole-brain afferent input mapping to functionally distinct brainstem noradrenaline cell types. Neurosci Res 194, 44–57 (2023).

54. Luskin, A. T., et al. A diverse network of pericoerulear neurons control arousal states. bioRxiv (2023) doi:10.1101/2022.06.30.498327.

55. Segalin, C. et al. The Mouse Action Recognition System (MARS) software pipeline for automated analysis of social behaviors in mice. Elife 10, e63720 (2021).

